# Tongue–Palate Electromyographic Synchronization Related to Swallowing, Mastication, and Speech

**DOI:** 10.1101/2025.07.17.665454

**Authors:** Hitoshi Maezawa, Kasane Kajimoto, Takuto Yoshimoto, Masanori Wakida

## Abstract

The specific oscillatory dynamics of intermuscular coupling involved in swallowing, mastication, speech production, and respiration have not been elucidated. This study aimed to explore intermuscular coupling between the tongue and palate that underlies oral functions by analyzing the event-related coherence (ERC) of electromyography (EMG) signals before swallowing, mastication, and speech production. Twenty-two healthy participants performed three oral tasks: swallowing, speech production, and mastication. Sixteen of these participants also completed a combined task involving swallowing immediately after speech. EMG signals were recorded from the tongue and palate, and ERC between tongue and palate EMG was analyzed. Peak ERC was compared across tasks for alpha (8–14 Hz), beta (16–30 Hz), and gamma (32–46 Hz) frequency bands. In all participants, ERC showed significant peaks in all frequency bands before the onset of swallowing, mastication, and speech. Only the ERC values in the beta band were significantly larger for swallowing than for mastication, but not for speech. Moreover, the ERC values for the combined task of swallowing after speech production were significantly smaller than those for the simple swallowing task in the alpha, beta, and gamma bands. The intermuscular oscillatory regulation is more critical for swallowing than for mastication.

## Introduction

The oral region is essential for various body functions, such as swallowing, mastication, speech production, and respiration. These functions are spatiotemporally regulated by the sophisticated and coordinated movements of multiple oropharyngeal and facial structures. In particular, intermuscular coordination between the tongue and palate is crucial for the precise control of these essential oral functions. For instance, at the onset of swallowing, the tongue rises to the palate, creating negative pressure in the oral cavity and inducing the swallowing reflex [1]. During speech, various vowels are produced by changing the contact point between the tongue and palate [2]. During mastication, the tongue and soft palate work in tandem to efficiently chew food and facilitate a smooth transition to swallowing [3]. Previous studies using electromyography (EMG) have highlighted the importance of neuromuscular functions in these processes, with particular attention to the role of the timing and amplitude of EMG activity during swallowing [4], mastication [5], and speech production [6]. However, little is known about the fine coordination between the tongue and palate in the neuromuscular control of these vital oral functions.

Numerous studies have investigated the descending neural drive from the primary sensorimotor cortex (SM1), primarily by analyzing the frequency domain of the coupling between brain and muscle activity during voluntary finger contractions [7–12]. The beta and gamma bands—which are closely related to the corticospinal drive from SM1 to the contralateral muscles—have been shown to exhibit evidence of corticomuscular coherence [13–15]. A previous study reported that tongue movements are regulated by the SM1 in both hemispheres via beta oscillation through the corticobulbar tract [16]. Another method often used to examine motor control characteristics is intermuscular coherence, which involves analyzing the linear dependencies between two EMG signals at a specific frequency [17]. This method allows for the non-invasive study of common synaptic inputs to motor neuron pools across muscles in humans [18]. Intermuscular coherence in the upper and lower limbs has been linked to cortical and spinal mechanisms [19–22]. The functional connection between muscles has been demonstrated using motor-relevant oscillatory components at alpha, beta, and gamma frequencies [23]. However, although intermuscular coupling has been detected in the upper and lower limbs, little is known about the specific oscillatory dynamics of intermuscular coupling in the oral region during the execution of oral functions. The fine coordination between oral structures, particularly the tongue and palate, ensures that each oral function is performed smoothly and safely, preventing complications such as aspiration or unclear speech. In this study, we aimed to investigate the event-related coherence (ERC) of EMG signals between the tongue and palate prior to swallowing, mastication, and speech production. By analyzing ERC across different frequency bands (alpha, beta, and gamma), we sought to elucidate the underlying oscillatory synchronization that facilitates these critical oral functions. Understanding the oscillatory synchronization between the tongue and palate could provide deeper insights into neuromuscular control and its role in various physiological and communicative processes, such as swallowing, speech production, and mastication.

We hypothesized that the peak ERC values would differ among swallowing, speech, and mastication, given the varying involvement of the tongue and palate depending on the oral function being performed. Furthermore, by comparing the coherence values for the combined task of swallowing immediately after speech with those for simple swallowing, we aimed to evaluate the mechanisms of intermuscular communication during the “eating while talking” situation, which often leads to aspiration in daily life.

## Results

Figure 1(a) shows 2-s epochs of raw rectified EMG signals from the tongue and palate, and rectified electroglottography signal for swallowing in a representative participant (participant 18). The EMG activity in the tongue and palate muscles is shown prior to the onset of swallowing (dashed line).

**Figure 1.**
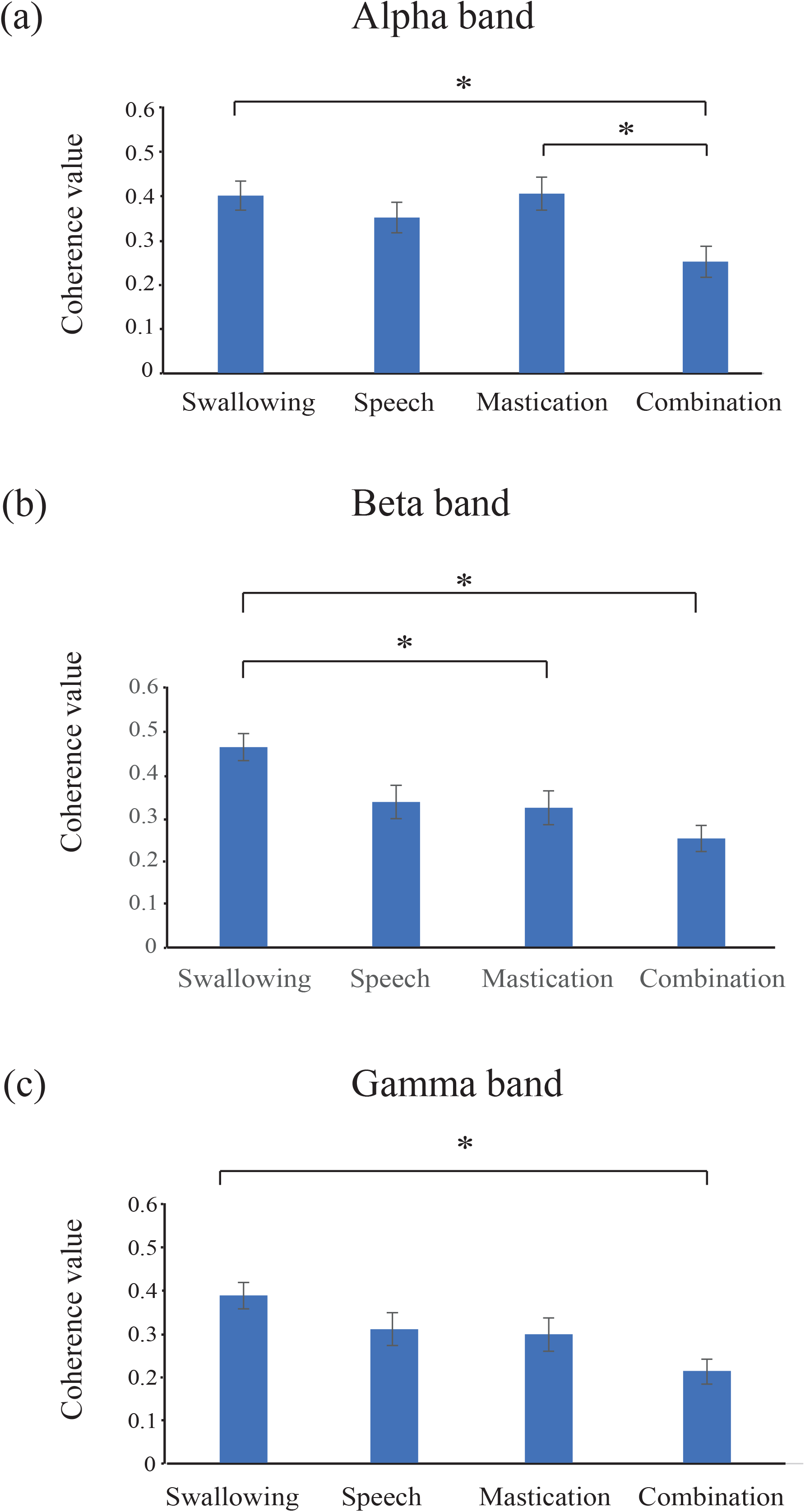
Raw signals of electroglottography and EMG **(a)** and time-frequency plots of coherence **(b)** for swallowing in a single participant (participant 18). (**a**) 2-s epochs of raw rectified electroglottography signals for swallowing. The vertical dashed line indicates the onset of electroglottography signals for swallowing. In addition, 2-s epochs of raw rectified EMG signals from the tongue and palate for swallowing are shown. The EMG signals are activated in both the tongue and palate before the start of swallowing. (**b**) Time-frequency plots of coherence values for swallowing. A significant coherence is observed between the tongue and palate in the alpha, beta, and gamma frequency bands prior to the onset of swallowing. The vertical line indicates the frequency range from 8 Hz to 46 Hz. The horizontal line indicates the time from 1.5 to 0 s before the onset of swallowing. EMG, electromyography; Alpha, Alpha frequency band; Beta, Beta frequency band; Gamma, Gamma frequency band.

Figure 1(b) shows time-frequency plots of the coherence values from swallowing for a single participant (participant 18). A significant ERC between the tongue and palate was observed in the alpha, beta, and gamma frequency bands prior to the onset of each task. Individual peak coherence values for the alpha, beta, and gamma bands in each task are presented in Tables 1, 2, and 3, respectively.

**Table 1.** Individual coherence values at the alpha band in each task.

| Par | Swallow |  | Speech |  | Mastication |  | Combination |  |
| --- | --- | --- | --- | --- | --- | --- | --- | --- |
|  | Freq (Hz) | Value | Freq (Hz) | Value | Freq (Hz) | Value | Freq (Hz) | Value |
| 1 | 12 | 0.248 | 8 | 0.200 | 12 | 0.299 |  |  |
| 2 | 14 | 0.235 | 10 | 0.175 | 10 | 0.245 |  |  |
| 3 | 10 | 0.105 | 10 | 0.230 | 14 | 0.197 |  |  |
| 4 | 10 | 0.323 | 8 | 0.162 | 10 | 0.545 |  |  |
| 5 | 12 | 0.363 | 10 | 0.297 | 8 | 0.384 |  |  |
| 6 | 8 | 0.348 | 8 | 0.596 | 8 | 0.296 |  |  |
| 7 | 12 | 0.508 | 10 | 0.270 | 10 | 0.210 | 14 | 0.071 |
| 8 | 14 | 0.432 | 14 | 0.353 | 14 | 0.489 | 14 | 0.266 |
| 9 | 12 | 0.352 | 14 | 0.226 | 10 | 0.449 | 8 | 0.133 |
| 10 | 10 | 0.493 | 12 | 0.603 | 12 | 0.144 | 14 | 0.274 |
| 11 | 10 | 0.265 | 12 | 0.328 | 12 | 0.283 | 14 | 0.194 |
| 12 | 8 | 0.646 | 12 | 0.278 | 8 | 0.684 | 10 | 0.221 |
| 13 | 14 | 0.788 | 14 | 0.275 | 14 | 0.481 | 8 | 0.320 |
| 14 | 10 | 0.540 | 8 | 0.617 | 8 | 0.539 | 8 | 0.536 |
| 15 | 14 | 0.321 | 8 | 0.218 | 8 | 0.441 | 10 | 0.403 |
| 16 | 14 | 0.359 | 10 | 0.309 | 10 | 0.189 | 10 | 0.245 |
| 17 | 8 | 0.509 | 8 | 0.252 | 12 | 0.224 | 14 | 0.294 |
| 18 | 10 | 0.351 | 8 | 0.417 | 12 | 0.648 | 10 | 0.472 |
| 19 | 14 | 0.456 | 12 | 0.632 | 14 | 0.720 | 14 | 0.144 |
| 20 | 8 | 0.526 | 10 | 0.222 | 12 | 0.314 | 12 | 0.067 |
| 21 | 8 | 0.306 | 14 | 0.622 | 14 | 0.554 | 14 | 0.297 |
| 22 | 8 | 0.271 | 8 | 0.385 | 8 | 0.508 | 10 | 0.054 |
| Ave | 10.9 | 0.3975 | 10.4 | 0.3485 | 10.9 | 0.4020 | 11.5 | 0.2494 |
| Min | 8 | 0.105 | 8 | 0.162 | 8 | 0.144 | 8 | 0.054 |
| Max | 14 | 0.788 | 14 | 0.632 | 14 | 0.720 | 14 | 0.536 |
| SEM | 0.5 | 0.0324 | 0.5 | 0.0324 | 0.5 | 0.0367 | 0.6 | 0.0352 |

Freq, frequency; Par, participant number; Value, peak value

**Table 2.**
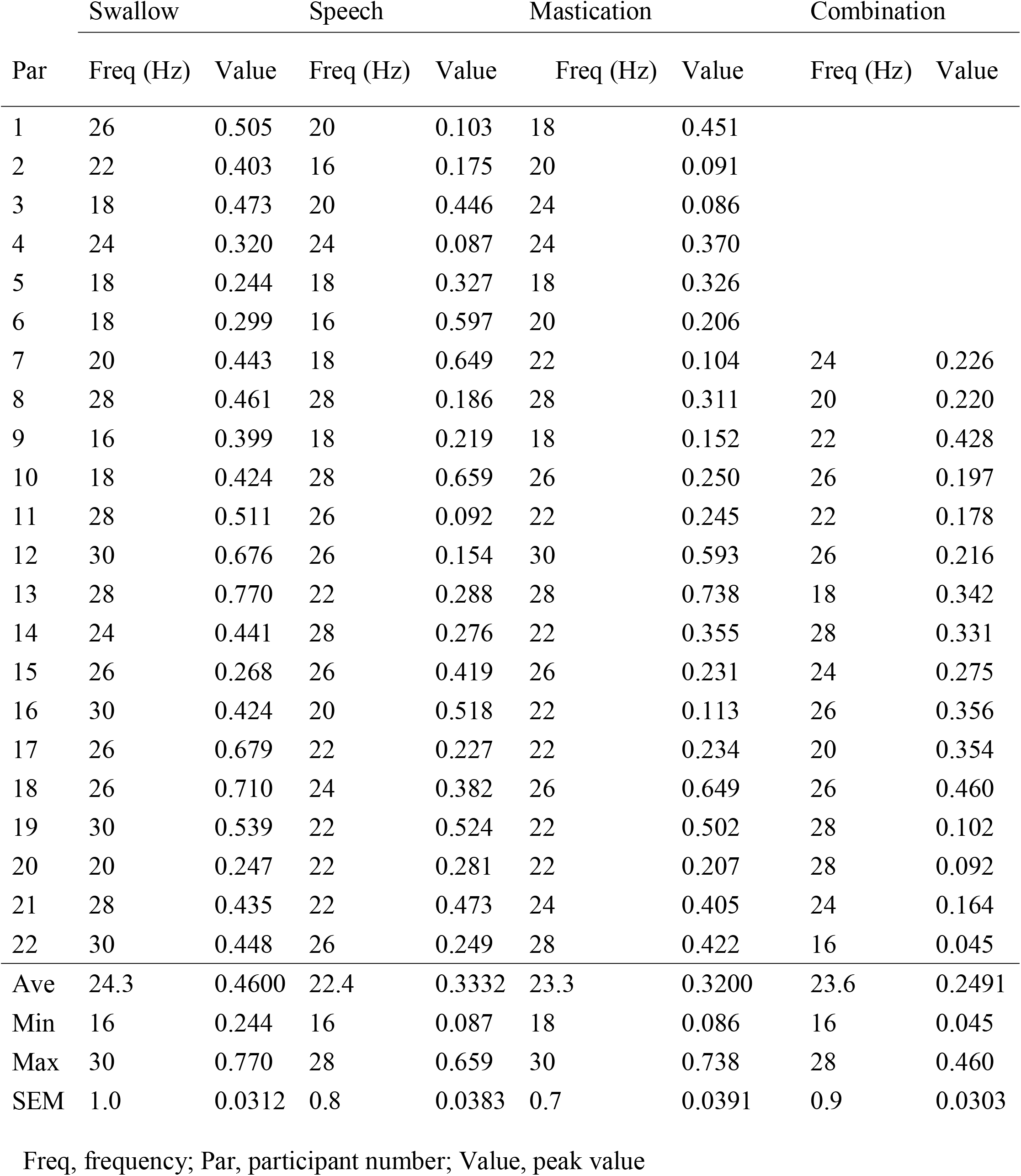
Individual coherence values at the beta band in each task.

| Par | Swallow |  | Speech |  | Mastication |  | Combination |  |
| --- | --- | --- | --- | --- | --- | --- | --- | --- |
|  | Freq (Hz) | Value | Freq (Hz) | Value | Freq (Hz) | Value | Freq (Hz) | Value |
| 1 | 26 | 0.505 | 20 | 0.103 | 18 | 0.451 |  |  |
| 2 | 22 | 0.403 | 16 | 0.175 | 20 | 0.091 |  |  |
| 3 | 18 | 0.473 | 20 | 0.446 | 24 | 0.086 |  |  |
| 4 | 24 | 0.320 | 24 | 0.087 | 24 | 0.370 |  |  |
| 5 | 18 | 0.244 | 18 | 0.327 | 18 | 0.326 |  |  |
| 6 | 18 | 0.299 | 16 | 0.597 | 20 | 0.206 |  |  |
| 7 | 20 | 0.443 | 18 | 0.649 | 22 | 0.104 | 24 | 0.226 |
| 8 | 28 | 0.461 | 28 | 0.186 | 28 | 0.311 | 20 | 0.220 |
| 9 | 16 | 0.399 | 18 | 0.219 | 18 | 0.152 | 22 | 0.428 |
| 10 | 18 | 0.424 | 28 | 0.659 | 26 | 0.250 | 26 | 0.197 |
| 11 | 28 | 0.511 | 26 | 0.092 | 22 | 0.245 | 22 | 0.178 |
| 12 | 30 | 0.676 | 26 | 0.154 | 30 | 0.593 | 26 | 0.216 |
| 13 | 28 | 0.770 | 22 | 0.288 | 28 | 0.738 | 18 | 0.342 |
| 14 | 24 | 0.441 | 28 | 0.276 | 22 | 0.355 | 28 | 0.331 |
| 15 | 26 | 0.268 | 26 | 0.419 | 26 | 0.231 | 24 | 0.275 |
| 16 | 30 | 0.424 | 20 | 0.518 | 22 | 0.113 | 26 | 0.356 |
| 17 | 26 | 0.679 | 22 | 0.227 | 22 | 0.234 | 20 | 0.354 |
| 18 | 26 | 0.710 | 24 | 0.382 | 26 | 0.649 | 26 | 0.460 |
| 19 | 30 | 0.539 | 22 | 0.524 | 22 | 0.502 | 28 | 0.102 |
| 20 | 20 | 0.247 | 22 | 0.281 | 22 | 0.207 | 28 | 0.092 |
| 21 | 28 | 0.435 | 22 | 0.473 | 24 | 0.405 | 24 | 0.164 |
| 22 | 30 | 0.448 | 26 | 0.249 | 28 | 0.422 | 16 | 0.045 |
| Ave | 24.3 | 0.4600 | 22.4 | 0.3332 | 23.3 | 0.3200 | 23.6 | 0.2491 |
| Min | 16 | 0.244 | 16 | 0.087 | 18 | 0.086 | 16 | 0.045 |
| Max | 30 | 0.770 | 28 | 0.659 | 30 | 0.738 | 28 | 0.460 |
| SEM | 1.0 | 0.0312 | 0.8 | 0.0383 | 0.7 | 0.0391 | 0.9 | 0.0303 |

Freq, frequency; Par, participant number; Value, peak value

**Table 3.** Individual coherence values at the gamma band in each task.

| Par | Swallow |  | Speech |  | Mastication |  | Combination |  |
| --- | --- | --- | --- | --- | --- | --- | --- | --- |
|  | Freq (Hz) | Value | Freq (Hz) | Value | Freq (Hz) | Value | Freq (Hz) | Value |
| 1 | 32 | 0.494 | 32 | 0.132 | 32 | 0.343 |  |  |
| 2 | 32 | 0.337 | 34 | 0.217 | 42 | 0.094 |  |  |
| 3 | 34 | 0.104 | 40 | 0.075 | 42 | 0.049 |  |  |
| 4 | 36 | 0.410 | 36 | 0.231 | 36 | 0.383 |  |  |
| 5 | 34 | 0.332 | 40 | 0.499 | 32 | 0.413 |  |  |
| 6 | 36 | 0.301 | 32 | 0.434 | 32 | 0.114 |  |  |
| 7 | 36 | 0.304 | 36 | 0.446 | 44 | 0.099 | 34 | 0.156 |
| 8 | 36 | 0.448 | 44 | 0.197 | 40 | 0.117 | 40 | 0.187 |
| 9 | 38 | 0.250 | 42 | 0.139 | 38 | 0.231 | 42 | 0.421 |
| 10 | 36 | 0.543 | 32 | 0.777 | 32 | 0.203 | 34 | 0.192 |
| 11 | 36 | 0.283 | 38 | 0.137 | 32 | 0.200 | 44 | 0.132 |
| 12 | 44 | 0.603 | 44 | 0.223 | 38 | 0.621 | 32 | 0.155 |
| 13 | 34 | 0.583 | 38 | 0.182 | 32 | 0.572 | 38 | 0.181 |
| 14 | 36 | 0.520 | 36 | 0.371 | 34 | 0.631 | 36 | 0.450 |
| 15 | 34 | 0.216 | 32 | 0.355 | 36 | 0.407 | 32 | 0.256 |
| 16 | 32 | 0.522 | 36 | 0.205 | 38 | 0.218 | 34 | 0.220 |
| 17 | 32 | 0.377 | 36 | 0.429 | 38 | 0.154 | 44 | 0.305 |
| 18 | 42 | 0.309 | 36 | 0.520 | 36 | 0.405 | 36 | 0.215 |
| 19 | 36 | 0.662 | 36 | 0.568 | 36 | 0.564 | 42 | 0.213 |
| 20 | 44 | 0.401 | 32 | 0.089 | 38 | 0.249 | 38 | 0.053 |
| 21 | 44 | 0.243 | 44 | 0.316 | 38 | 0.229 | 44 | 0.243 |
| 22 | 32 | 0.311 | 32 | 0.294 | 32 | 0.273 | 32 | 0.034 |
| Ave | 36.2 | 0.3888 | 36.7 | 0.3107 | 36.3 | 0.2986 | 37.6 | 0.2133 |
| Min | 32 | 0.104 | 32 | 0.075 | 32 | 0.049 | 32 | 0.034 |
| Max | 44 | 0.662 | 44 | 0.777 | 44 | 0.631 | 44 | 0.450 |
| SEM | 0.8 | 0.0304 | 0.9 | 0.0383 | 0.8 | 0.0382 | 1.1 | 0.0277 |

Freq, frequency; Par, participant number; Value, peak value

Analysis of variance revealed a main effect of the task in the alpha (F = 6.105, df = 3, *P* = 0.001), beta (F = 6.758, df = 1.963, *P* = 0.004), and gamma (F = 5.085, df = 3, *P* = 0.004) frequency bands (Tables 1, 2, and 3, respectively). Post-hoc analyses revealed that the ERC values for swallowing and mastication were significantly higher than that for the combined task of swallowing after speech in the alpha frequency band [Figure 2(a)]. In addition, the ERC value for swallowing was significantly higher than that for mastication and combined task of swallowing after speech in the beta frequency band [Figure 2(b)]. Furthermore, the ERC value for swallowing was significantly higher than that for the combined task of swallowing after speech in the gamma frequency band [Figure 2(c)].

**Figure 2.**
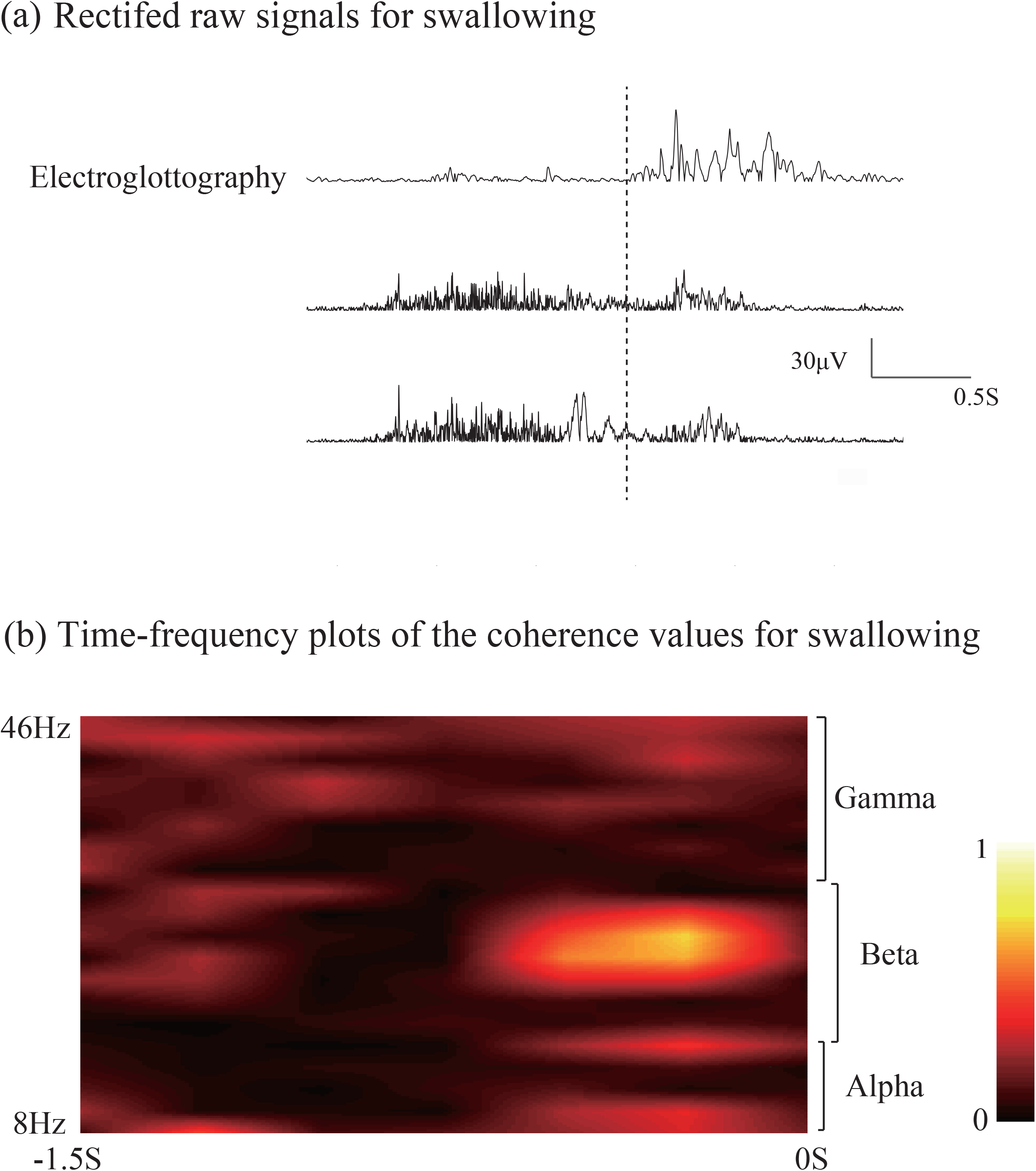
Mean values of event-related coherence at the alpha **(a)**, beta **(b)**, and gamma **(c)** bands across participants. (**a**) Coherence values for swallowing and mastication are significantly larger than those for the combined task of swallowing after speech in the alpha band. (**b**) The coherence value for swallowing is significantly larger than that for mastication and the combined task of swallowing after speech in the beta band. (**c**) The coherence value for swallowing is significantly larger than that for the combined task of swallowing after speech in the gamma band. Combination, Combined task of swallowing after speech.

## Discussion

In the present study, we investigated the ERC of EMG signals between the tongue and palate prior to swallowing, mastication, and speech production. The results showed that intermuscular coherence between the tongue and palate was detected prior to the onset of vital oral functions, such as swallowing, speech, and mastication. The coherence was stronger in the beta band for swallowing than for mastication, but not for speech. Moreover, coherence during the combined task of swallowing after speech was significantly weaker than that during simple swallowing in the alpha, beta, and gamma frequency bands.

Significant ERC was observed in the alpha, beta, and gamma frequency bands prior to the onset of swallowing, speech, and mastication, suggesting that these oral tasks are regulated by fine-tuned oscillatory interactions between the tongue and palate across different oscillatory frequency bands. In addition, ERC values in the beta band, but not in the alpha and gamma bands, were higher for swallowing than for mastication, but not for speech. It is known that both beta and gamma intermuscular coherence mainly reflect the corticomuscular drive from the SM1 to peripheral regions through motor neuron pools [14, 19, 24], and that coherence in the beta and gamma frequency bands between the tongue and palate mainly reflects efferent-related oscillatory synchronization between them [16]. Thus, the stronger ERC in the beta frequency band for swallowing than for mastication may suggest that efferent-related fine-tuning between the tongue and palate, regulated by SM1, is particularly important for swallowing than for mastication.

During swallowing, precise coordination between the tongue and palate is critical for the safe transportation of food and liquids from the oral cavity to the pharynx and then to the esophagus. Swallowing begins with tongue elevation against the palate, creating negative pressure in the oral cavity, which is crucial for triggering the swallowing reflex [25]. This reflex generates a force that propels the food bolus from the posterior end of the oral cavity into the pharynx. Moreover, the contact between the tongue and palate temporarily seals the anterior portion of the oral cavity, preventing leakage of contents into the nasal cavity or the front end of the oral cavity. For these reasons, coordination between the tongue and palate is particularly critical for initiating swallowing.

In this study, a significant ERC value was detected for mastication; however, it was lower than that for swallowing in the beta band. Mastication is mainly performed by the main muscles of mastication, such as the masseter and temporalis, while the tongue and soft palate play a relatively smaller role in mastication than in swallowing [26]. This reduced involvement of the tongue and palate may explain the significantly lower coupling between them during mastication than during swallowing.

A significant ERC value was also detected for speech. Both swallowing and speech require critically coordinated movements of the tongue and palate [6]. Speech particularly necessitates a greater variety of tongue and palate movements to enable advanced lingual communication. Speech utilizes exhaled air from respiration to vibrate the vocal cords in the larynx, creating primary laryngeal sounds. These sounds then resonate in the oral, pharyngeal, and nasal cavities to produce vowels. The articulatory organs (i.e., the tongue, soft palate, and lips) move to narrow the vocal tract and generate noise, producing consonants. The pronunciation of a vowel is determined by the extent of soft palate elevation, the position of the tongue, and the degree to which the lips are open

In the study, the simple speech task involved pronouncing the syllable “Ta,” which requires the movement of both the tongue and the palate. To produce the consonant “T” in “Ta,” the anterior part of the tongue touches the alveolar ridge to momentarily stop the airflow. Then, the tongue quickly moves away to release the air, producing the “T” sound. After that, the tongue drops to the position for producing the vowel “a,” while the soft palate remains raised. The significant ERC observed between the tongue and palate suggests that coordination between both organs is crucial for speech production.

Interestingly, we observed a significantly decreased coherence for the combined task of swallowing immediately after speaking compared with that for simple swallowing in all the frequency bands (alpha, beta, and gamma bands). Both swallowing and speaking require coordinated movement of oropharyngeal structures, including the tongue and palate. However, the mechanisms of swallowing and speaking differ, and it may be occasionally difficult to perform these actions consecutively or simultaneously for several reasons [27]. Swallowing and speaking involve distinct muscle movements in widely distributed oropharyngeal and facial regions. During swallowing, the tongue moves food from the mouth into the pharynx, and the soft palate rises to shut off the nasal cavity. In contrast, speaking requires the tongue and palate to move to various positions to produce sounds. When these actions are performed simultaneously, these movements can compete, which may lead to incorrect swallowing. Additionally, when talking while eating, frequent switching between swallowing and speaking is required. If this rapid switch is disrupted, the swallowing reflex may become incomplete, thereby increasing the likelihood of aspiration. When speaking and swallowing consecutively, the oral preparation period is shorter than during simple swallowing, and coordinated movements may not be sufficiently effective. Thus, the observed decrease in coherence strength during the combined tasks of swallowing immediately after speech may be related to insufficient coordination between the tongue and palate. It is advisable to give due care to this issue in daily life to reduce the risk of aspiration resulting from speaking while eating [27].

Our study has some limitations. Speech in daily life is not limited to single syllables but involves a more complex combination of vowels and consonants to form words and sentences. In this study, the task of speech production was limited to producing the single sound “Ta.” Other vowels and consonants are likely to exhibit different forms of coordination between the tongue and palate, which may affect ERC strength. In addition, the evaluation of the mastication task in this study did not involve the insertion of food into the mouth. During eating, the tongue plays a role in manipulating food within the oral cavity, positioning it optimally for mastication by the teeth. Therefore, the tongue movements observed in the study diverged from the complex patterns typically associated with mastication of food. Further studies are required to investigate the ERC between the tongue and palate during speech production tasks that approximate conversations using multiple words and mastication tasks involving the chewing of food.

In conclusion, our findings demonstrate that essential oral functions, including swallowing, speech, and mastication, are intricately regulated by intermuscular coupling between the tongue and palate that is evident in the alpha, beta, and gamma frequency bands. The regulation of intermuscular coupling was proven to be particularly important for swallowing; however, it was reduced during the combined task of swallowing immediately after speech production, indicating insufficient intermuscular communication between the tongue and palate during the situation of “eating while talking.” This situation, commonly seen in daily life, may lead to aspiration and choking.

## Methods

### Participants

Twenty-two healthy volunteers (15 men; age range, 20–25 years; mean age, 21.1 years) were examined. All participants were right-handed, and none had a history of neurological or psychiatric disorders.

The study was conducted in accordance with the ethical standards stated in the Declaration of Helsinki and approved by the local Ethics and Safety Committees at Kansai Medical University (No. 2020107). All participants provided written informed consent prior to enrolment in the study.

### Recording the Oral Tasks

All participants performed three experimental tasks: swallowing, speech, and mastication. In addition, 16 participants were asked to perform a combined task that included swallowing after speech. The order of the three (or four) tasks was randomized for each participant, with several minutes of rest between tasks. During the experimental tasks, participants were comfortably seated in a chair and instructed to fix their gaze on a point on the wall to avoid any effects from eye movement or visual perception.

For the swallowing task, participants were asked to perform repetitive, volitional swallowing of their saliva at their own pace in intervals of approximately 8 s for at least 8 min. For the speech production task, participants were asked to repetitively produce the single syllable “Ta” at their own pace, with intervals of approximately 5 s for at least 5 min. For the mastication task, participants were also asked to participate in a single mastication session at their own pace, with intervals of approximately 5 s for at least 5 min. For the combined task, which included swallowing after speech, participants were asked to continuously produce speech with the three syllables “Pa,” “Ta,” and “Ka,” and then to swallow their saliva at their own pace, with intervals of approximately 8 s for at least 8 min.

Bipolar surface EMGs were recorded bilaterally from the dorsum of the tongue and the anterior part of the soft palate using disposable electrodes (15×15 mm; Vitrode V; Nihon Kohden, Tokyo, Japan) placed 30 mm apart [16, 28, 29]. Electroglottography signals were recorded from the epiglottis to examine the onset of swallowing and speech. To examine the onset of mastication, bipolar surface EMGs were recorded on both sides of the masticatory muscles using a pair of Ag/AgCl disk electrodes. EMG and electroglottography signals were sampled at 2000 Hz using a wireless EMG acquisition system (Ultium EMG; Noraxon USA Inc., Scottsdale, AZ, USA).

### Data Analysis

For coherence analysis, rectified raw EMG data from the tongue and palate were bandpass-filtered at 1–100 Hz. The onset of each trial was determined by the beginning of the electroglottography/EMG bursts, which were visually identified from the continuously recorded data offline. Rectified electroglottography was used for swallowing and speech production, while rectified EMG from the masticatory muscles was used for mastication. Each burst of the electroglottography and EMG signals was visually reviewed, and epochs containing ambiguous electroglottography and/or EMG bursts associated with unintended oropharyngeal movements were omitted from the analysis. The average number of trials per participant was 58.2, 57.9, 58.5, and 58.3 for the swallowing, mastication, speech production, and swallowing-after-speech tasks, respectively.

The coherence spectra of the rectified EMG signals between the tongue and palate were calculated using Welch’s method [30] of spectral density estimation with a Hanning window (2 Hz frequency resolution, 250 ms time resolution). The center of the moving window was shifted from 2 s before to 1 s after the movement onset. Coherence values (Cohxy) were calculated using the following equation:

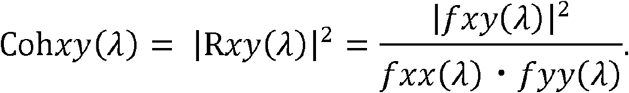

In this equation, *fxx(λ)* and *fyy(λ)* are the auto spectra values of the EMG signals of the tongue and palate for a given frequency *λ*, respectively, and *fxy(λ)* is the cross spectrum between them. Coherence is expressed as a real number between 0 and 1, where 1 indicates a perfect linear association between two signals and 0 indicates a complete absence of linear association.

Frequency bands of 8–14 Hz (alpha), 16–30 Hz (beta), and 32–46 Hz (gamma) were analyzed. For each frequency band, the peak value was determined in the band showing the maximal coherence value between 1.5 s and 0.25 s before the onset of each task. According to the method proposed by Rosenberg et al. [31], all coherence values above Z were considered significant at P < 0.05, where Z = 1 − 0.05(1/L − 1), and L denotes the total number of samples for the analysis of auto- and cross-spectra.

### Statistical Analyses

The data are expressed as mean ± standard error of the mean. Coherence values were normalized using an arc hyperbolic tangent transformation to stabilize variance [32]. The coherence values in each frequency band were then entered as dependent variables into a one-way repeated measures analysis of variance, with the following within-participant factor: tasks—swallowing, speech, mastication, and the combined task including swallowing after speech. After performing Mauchly’s sphericity test, we applied the Greenhouse–Geisser correction where necessary. When significant effects were identified by the analysis of variance, we subsequently employed exploratory post-hoc Student’s paired t-tests (two-tailed; Bonferroni correction for multiple comparisons) between each pair of tasks (swallowing, speech, mastication, and swallowing after speech) in each frequency band. The level of statistical significance was set at *P* < 0.05. All statistical analyses were performed using SPSS software (IBM Corp., Armonk, NY, USA).

## Data availability statement

Data presented in this study will be made available upon reasonable request and with permission of the study participants and a formal data sharing agreement. The data are available on request from the corresponding author.

## Acknowledgements

This work was supported by Grants-in-Aid for Scientific Research from the Japan Society for the Promotion of Science [grant numbers (B) 22H03452 (HM)].

## Author contributions

H. M. Contributed to conception, design, data acquisition and interpretation, performed all statistical analyses, drafted, and critically revised the manuscript. K. K. Contributed to conception, design, and critically revised the manuscript. T. Y. Contributed to conception, design, and critically revised the manuscript. M. W.: Contributed to conception, design, and critically revised the manuscript. All authors gave their final approval and agreed to be accountable for all aspects of the work.

## Additional Information

<u>Competing Interests Statement</u>

## References

1. Pitts, T., & Iceman, K. E. Deglutition and the regulation of the swallow motor pattern. Physiol. (Bethesda) 38, 10–24 (2023).

2. Gibbon, F. E., Lee, A. & Yuen, I. Tongue-palate contact during selected vowels in normal speech. Cleft Palate Craniofac. J. 47, 405–412 (2010).

3. Matsuo, K. & Palmer, J. B. Anatomy and physiology of feeding and swallowing: normal and abnormal. Phys. Med. Rehabil. Clin. N. Am. 19, 691–707 (2008).

4. Vaiman, M., Eviatar, E. & Segal, S. Surface electromyographic studies of swallowing in normal subjects: a review of 440 adults. Report 2. Quantitative data: amplitude measures. Report 2. Otolaryngol. Head Neck Surg. 131, 773–780 (2004).

5. Wood, W. W. A review of masticatory muscle function. J. Prosthet. Dent. 57, 222–232 (1987).

6. Balata, P. M., Silva, H. J., Moraes, K. J., Pernambuco, A. & Moraes, S. R. Use of surface electromyography in phonation studies: an integrative review. Int. Arch. Otorhinolaryngol. 17, 329–339 (2013).

7. Hansen, S., Hansen, N. L., Christensen, L. O., Petersen, N. T. & Nielsen, J. B. Coupling of antagonistic ankle muscles during co-contraction in humans. Exp. Brain Res. 146, 282–292 (2002).

8. Halliday, D. M. et al. Functional coupling of motor units is modulated during walking in human subjects. J. Neurophysiol. 89, 960–968 (2003).

9. Norton, J. A. & Gorassini, M. A. Changes in cortically related intermuscular coherence accompanying improvements in locomotor skills in incomplete spinal cord injury. J. Neurophysiol. 95, 2580–2589 (2006).

10. Nielsen, J. B. et al. Reduction of common motoneuronal drive on the affected side during walking in hemiplegic stroke patients. Clin. Neurophysiol. 119, 2813–2818 (2008).

11. Petersen, N. T. et al. Suppression of EMG activity by transcranial magnetic stimulation in human subjects during walking. J. Physiol. 537, 651–656 (2001).

12. Willerslev-Olsen, M., Petersen, T. H., Farmer, S. F. & Nielsen, J. B. Gait training facilitates central drive to ankle dorsiflexors in children with cerebral palsy. Brain 138, 589–603 (2015).

13. Conway, B. A. et al. Synchronization between motor cortex and spinal motoneuronal pool during the performance of a maintained motor task in man. J. Physiol. 489, 917–924 (1995).

14. Mima, T. & Hallett, M. Corticomuscular coherence: a review. J. Clin. Neurophysiol. 16, 501–511 (1999).

15. Grosse, P., Cassidy, M. J. & Brown, P. EEG-EMG, MEG-EMG and EMG-EMG frequency analysis: physiological principles and clinical applications. Clin. Neurophysiol. 113, 1523–1531 (2002).

16. Maezawa, H. et al. Contralateral dominance of corticomuscular coherence for both sides of the tongue during human tongue protrusion: an MEG study. Neuroimage 101, 245–255 (2014).

17. Gross, J. et al. Cortico-muscular synchronization during isometric muscle contraction in humans as revealed by magnetoencephalography. J. Physiol. 527, 623–631 (2000).

18. Dideriksen, J. L. et al. Coherence of the surface EMG and common synaptic input to motor neurons. Front. Hum. Neurosci. 12, 207; 10.3389/fnhum.2018.00207 (2018)

19. Boonstra, T. W. The potential of corticomuscular and intermuscular coherence for research on human motor control. Front. Hum. Neurosci. 7, 855; 10.3389/fnhum.2013.00855 (2013)

20. Boonstra, T. W. & Breakspear, M. Neural mechanisms of intermuscular coherence: implications for the rectification of surface electromyography. J. Neurophysiol. 107, 796–807 (2012). 10.1152/jn.00066.2011, Pubmed:22072508.

21. Grosse, P., Kühn, A., Cordivari, C. & Brown, P. Coherence analysis in the myoclonus of corticobasal degeneration. Mov. Disord. 18, 1345–1350 (2003).

22. Yamanaka, E., Horiuchi, Y. & Nojima, I. EMG-EMG coherence during voluntary control of human standing tasks: a systematic scoping review. Front. Neurosci. 17, 1145751 (2023).

23. Laine, C. M. & Valero-Cuevas, F. J. Intermuscular coherence reflects functional coordination. J. Neurophysiol. 118, 1775–1783 (2017).

24. Farmer, S. F., Bremner, F. D., Halliday, D. M., Rosenberg, J. R. & Stephens, J. A. The frequency content of common synaptic inputs to motoneurons studied during voluntary isometric contraction in man. J. Physiol. 470, 127–155 (1993).

25. Kieser, J. A. et al. The role of oral soft tissues in swallowing function: what can tongue pressure tell us? Aust. Dent. J. 59(Suppl 1), 155–161 (2014).

26. Thexton, A. J. Mastication and swallowing: an overview. Br. Dent. J. 173, 197–206 (1992).

27. Roy, N., Stemple, J., Merrill, R. M. & Thomas, L. Dysphagia in the elderly: preliminary evidence of prevalence, risk factors, and socioemotional effects. Ann. Otol. Rhinol. Laryngol. 116, 858–865 (2007).

28. Maezawa, H. et al. Cortico-muscular synchronization by proprioceptive afferents from the tongue muscles during isometric tongue protrusion. Neuroimage 128, 284–292 (2016).

29. Maezawa, H. et al. Effects of bilateral anodal transcranial direct current stimulation over the tongue primary motor cortex on cortical excitability of the tongue and tongue motor functions. Brain Stimul. 13, 270–272 (2020).

30. Welch, P. The use of fast Fourier transform for the estimation of power spectra: a method based on time averaging over short, modified periodograms. IEEE Trans. Audio Electroacoust. (Trans Electroacous, A.) 15, 70–73 (1967).

31. Rosenberg, J. R., Amjad, A. M., Breeze, P., Brillinger, D. R. & Halliday, D. M. The Fourier approach to the identification of functional coupling between neuronal spike trains. Prog. Biophys. Mol. Biol. 53, 1–31 (1989).

32. Halliday, D. M. et al. A framework for the analysis of mixed time series/point process data—theory and application to the study of physiological tremor, single unit discharges and electromyogram. Prog. Biophys. Mol. Biol. 64, 237–278 (1995).

